# SEM^2^: A computational framework to model multiscale mechanics with subcellular elements

**DOI:** 10.1101/2023.07.07.548118

**Authors:** Sandipan Chattaraj, Michele Torre, Constanze Kalcher, Alexander Stukowski, Simone Morganti, Alessandro Reali, Francesco Silvio Pasqualini

## Abstract

Modeling multiscale mechanics in shape-shifting biological tissues in embryos, traditional, or engineered cell culture platforms (organoids, organs-on-chips) is both important and challenging. In fact, it is difficult to model relevant tissue-level structural changes mediated by discrete events at the cellular and subcellular levels, such as migration and proliferation. To accomplish this, we leveraged the subcellular element modeling (SEM) method, where ensembles of coarse-grained particles interacting via empirically defined potentials are used to model individual cells while preserving cell rheology. However, an explicit treatment of multiscale mechanics in SEM was missing. Here, we introduced SEM^2^, an extended version of the open-source software SEM++ and LAMMPS, enabling new analyses and visualization of particle-level stress and strain. We demonstrated various functionalities of SEM^2^ by simulating cell creep, migration, and proliferation in scenarios that recapitulate classical and engineered cell culture platforms. For every scenario, we highlight key mechanobiology that emerges spontaneously from particle interactions and discuss recent experimental evidence as qualitative validations of our simulations. The code for SEM2 is available on GitHub at https://github.com/Synthetic-Physiology-Lab/sem2.

## Introduction

Morphogenesis, the process by which tissues and organs change shape to acquire novel structures and functions during embryonic development, has long fascinated life scientists^1,2^ and physicists^3^. More recently, engineers became interested in how tissues build themselves to design better preclinical models, including organoids^1,4,5^ and organs-on-chips^6–8^. New tools in molecular biology^9,10^ and optical engineering^11–13^ have shown us that mechanical forces across spatial scales modulate morphogenesis^14^. Yet, we have a limited ability to computationally model these interactions^15^.

From a modeling perspective, morphogenetic events are characterized by complex rearrangements of tissue shape driven by the proliferation and migration of millions of cells^2,9,16^. In other words, we must model large non-linear deformations driven by discrete, local events in a continuously growing system. To model these events, continuous approaches treat developing tissues as deforming solids, flowing liquids and/or visco-poro-elastic combinations^17^. Alternatively, cells can be modeled as discrete objects interacting on a lattice^15,18,19^, or freely in space^3,20^. All modalities face tradeoffs. We can simulate cell mechanics with continuous models, but it is hard to incorporate discrete biology. Discrete models more naturally capture biological processes but are costly, and cell mechanics is lost (e.g., Potts models^19^) or limited to cell membranes (e.g., vertex modeling^3^). Yet, the mechanical forces at play in the intracellular space, or cytoplasm, can influence the mechanical behavior of entire tissues^21,22^.

To study subcellular mechanics during morphogenesis, we propose an extension to the class of methods known as subcellular element modeling (SEM) that incorporates mechanics: we call this approach SEM^2^. SEM is a discrete, off-lattice framework that treats cells as mechanically-competent agents^23,24^ capable of biologically relevant behaviors, such as flowing in blood flow^25^, migration^26^, proliferation^27,28^, and epithelia shaping^29,30^. In SEM, each cell is formed by *N*_*p*_ particles that provide a coarse-grained representation of the subcellular space (Fig. 1A). Two key results have made SEM relevant to mechanobiology. First, Thea Newman established the link with cell mechanics using experimentally measured cell rheology to define mildly attractive and mildly repulsive potentials that enforce adhesion and volume exclusion (Fig. 1B)^23,24^. Later, Petros Koumoutsakos introduced SEM++^27^, a C++ implementation of SEM that leverages the open-source and strongly supported LAMMPS library^31^ to speed up computations. Further, algorithmic manipulations of the particles have been used to model cell migration and proliferation^27^, while multiple particle types were introduced to model cell nuclei or membranes, explicitly^27,32^. Together these results have made SEM the framework of choice for modeling cell shape as an emergent property of subcellular interactions^33^. Yet, a treatment of multiscale mechanics in SEM is missing.

**Figure 1:**
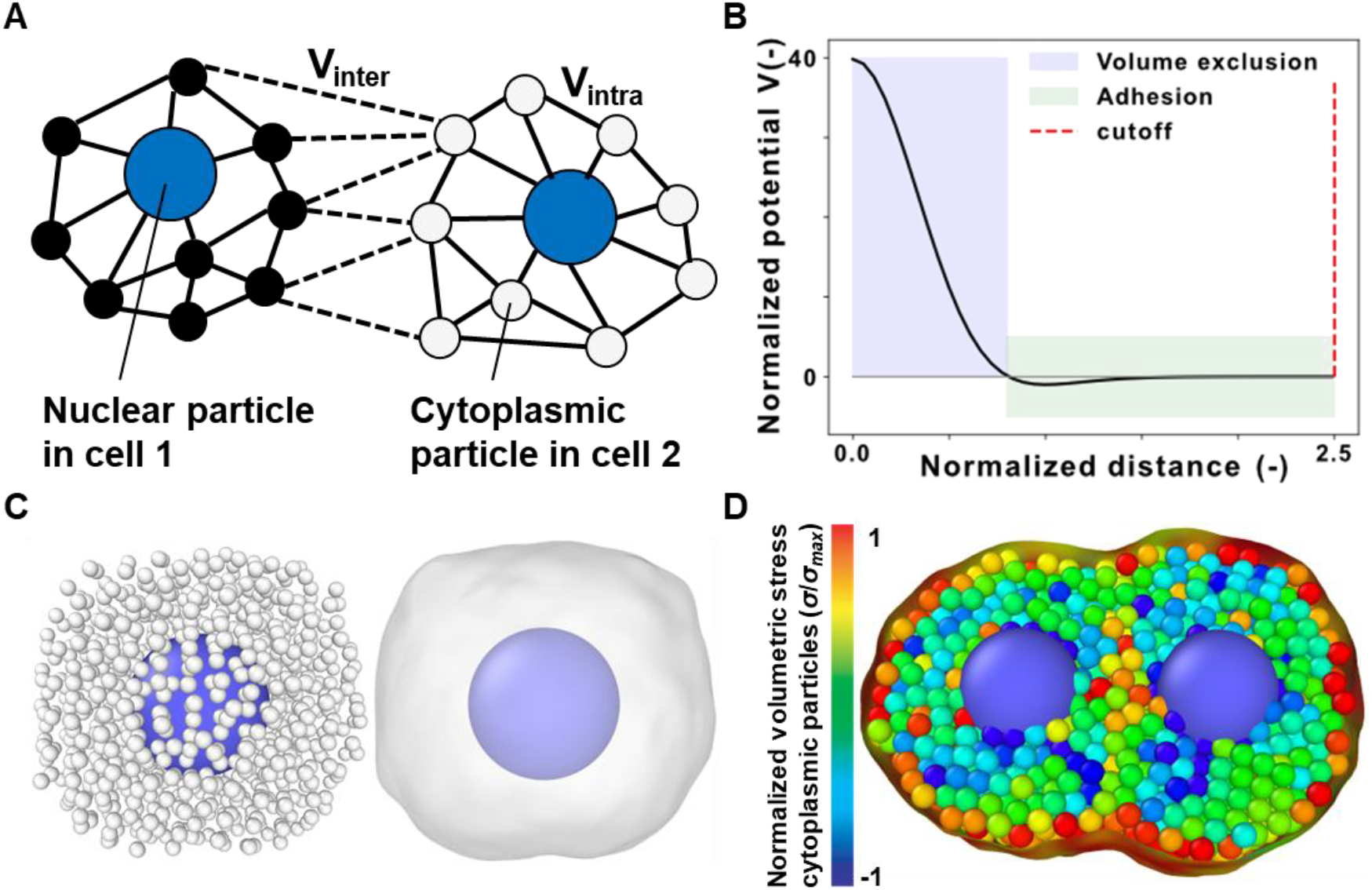
Subcellular element modeling and mechanics (SEM^2^). (A) A particle-based representation of two cells where the solid lines and dashed lines denote subcellular and intercellular interactions, respectively. (B) The plot of intracellular potential as a function of interparticle distance normalized to the equilibrium distance (potential minimum). (C) Particle-based and surface-based representations of a single cell with its nucleus. (D) A representation of a dividing cell in which the cytoplasmic particles are color-coded based on the per-particle volumetric stress (see Methods).

In SEM^2^, we realized that known potentials and computed particle displacement could be used to estimate subcellular mechanics with per-particle stress and strain and relate them to cell and tissue level deformation during morphogenesis. In this paper, we updated SEM++ and combined it with new analyses and visualizations to simulate particle ensembles (Fig. 1C) during key cellular behaviors, such as cell division (Fig. 1D), and study how stress and strain propagate across spatial scales. To demonstrate this approach, we present *in silico* studies of cells undergoing creep, migration, and proliferation in traditional and engineered cell culture environments. SEM^2^ and the input scripts for all presented simulations are available on GitHub (https://github.com/Synthetic-Physiology-Lab/sem2).

### SEM^2^ scalably recapitulates cell rheology in creep experiments

Simulating cells and tissues with SEM implies solving the Langevin equation for an arbitrary number of subcellular particles (*N*_*p*_)^23^.

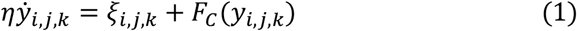

In Eq. (1), *η*is the viscous drag coefficient and *Y*_*i,j,k*_, 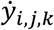are the position and velocity of each particle. We chose the indexes *i, j, k* to denote particle number, particle type, and cell number, respectively. That is, if we assume two cells are modeled using a single nuclear particle and many cytoplasmic ones per cell (two types, Fig. 1A), the positions of the 100^th^ cytoplasmic particle in cell 1 and the 300^th^ cytoplasmic particle in cell 2 would be denoted as *Y*_100,1,1_ and *Y*_300,1,2_. With this notation, *ξ*_*i,j,k*_ is the thermal fluctuation felt by each particle and *F*_*c*_(*Y*_*i,j,k*_) is the net pairwise force acting on it. In particular, Newman used the following empirically defined, mild potential (*V*(*d*) where *d* is the inter-particle distance) to enforce adhesion and volume exclusion (Fig 1b and Eq. 2)^23,27^.

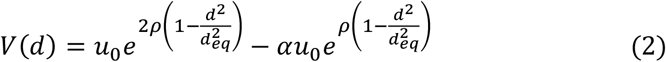

In Eq. (2), *u*_0_ is the potential’s well depth, *ρ* and *α* are scaling and shifting factors, and *d*_*eq*_ is the equilibrium distance between particles. To limit the computational cost, these potentials focus on short-range interactions and cut off at 2.5 times the equilibrium distance. Unless otherwise noted, we used parameter values as in SEM++^23^. To assign the forces in Eq. (1) based on cell-level rheology, Newman assumed *N*_*p*_ densely packed particles connected by springs and recovered the following relationships^23^

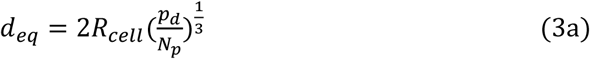

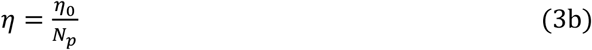

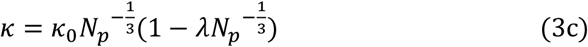

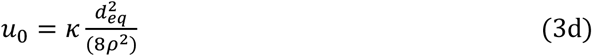

In Eqs. (3a-3d), *p*_*d*_ is the sphere close packing density. *R*_*cell*_, *κ*_0_, *η*_0_ are the radius, stiffness and viscosity of a cell, respectively; finally, *λ* = 0.75 is a tuning coefficient^23,27^.

In SEM^2^, we wanted to verify that these relations hold for arbitrary *N*_*p*_. To demonstrate it, we resorted to modeling a classical biomechanics experiment, cell creep. In this setup, cells are loaded between two glass pipettes attached to force transducers^34,35^. As one pipette is kept stationary, the other is displaced, thus applying controlled stress on the cell and generating measurable strain via cell lengthening (Fig. 2A). To simulate this experiment, we set up a simulation with a constrained cell composed of *N*_*p*_ particles, with *N*_*p*_ ranging from 250 to 10,000. To repeat these simulations as a function of particle number, one must convert the constant stress applied via the movable pipette in all simulations to the appropriate external force experienced by the particles in contact with the glass pipettes. However, the surface area in contact with the glass pipette and the precise number of particles in that area vary nonlinearly with *N*_*p*_ based on cell geometry and particle stacking (Supplementary Fig. S1). To solve this problem, we first created two parallel, flat surfaces, known as *slabs* in LAMMPS (black rectangle, Fig. 2B)^23,24^. Then, we calculated the surface area (red shared area, Fig. 2B) of the slabs and used it to convert the applied stress to the total force that we allocated to the particles in the top slab (See Methods).

**Figure 2:**
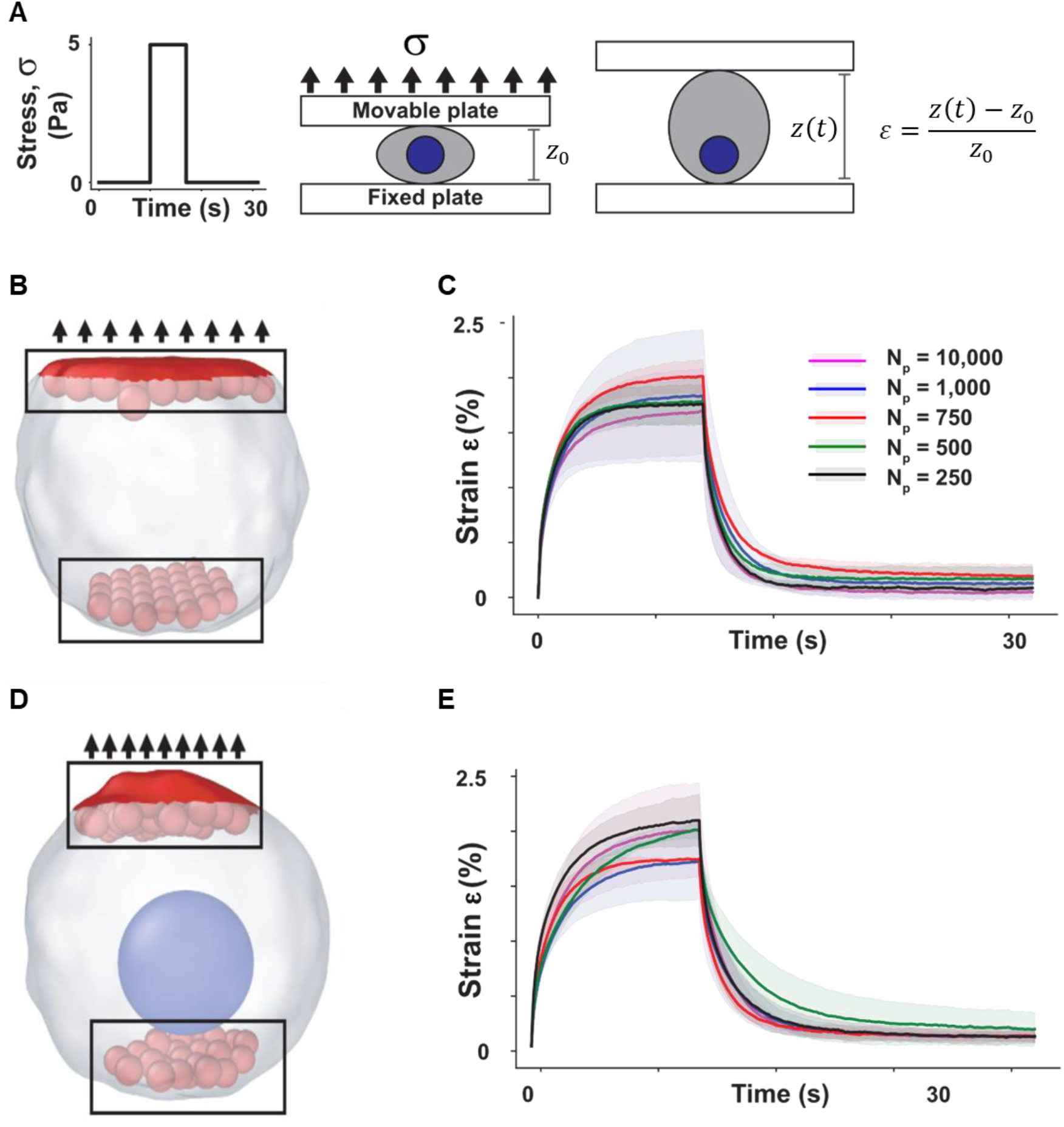
SEM^2^ scalably captures cell rheology. (A) Illustration of a single-cell creep experiment. Constant stress (5 Pa) is applied to a cell sandwiched between a fixed and a movable plate, resulting in an axial strain. (B) The cell in the pre-stretched configuration is plotted using a Gaussian surface mesh (white) that encloses all particles (see also 1C and Methods). The top red (shaded) surface indicates the area considered for stress calculation. The black rectangles encompass the particles (shown in red) in the fixed and movable slabs. (C) Creep strain percentage versus time for a single cell with a single particle type. Solid lines indicate mean time courses across three different simulation runs, shaded areas are standard deviations. (D) The cell (white) with a single nuclear particle (blue) in the pre-stretched configuration, as in Fig 2B. (E) Creep strain percentage versus time for a single cell with cytoplasmic and nuclear particles, as in Fig. 2C.

Having standardized force application, we asked whether the relationships in Eqs. (3a-3d) could be regarded as scaling laws: that is, do the cells composed of 250 and 10,000 particles undergo the same strain, under the same applied stress? To answer this question, we first considered a cell composed of a single particle type (no nucleus, Fig. 2B) and subjected it to a short-duration stress (5 Pa for 7 s, Fig 2A). To account for the effect of stochastic thermal fluctuations, we performed three different simulations per value of *N*_*p*_ and plotted the average (solid curve) and the standard deviation (shaded area, Fig 2C) for each condition. Our results show that SEM^2^ predicts a similar strain time course for all simulated *N*_*p*_ (Fig. 2C). Second, following Milde *et al*.,^27^ we introduced the cell nucleus as a more massive particle of greater radius (Fig. 2D, Supplementary Video SV1) which is accomplished by using a shifted version of the potential in Eq. (2) to accommodate the nuclear radius. Then, we repeated the simulations to demonstrate that the scalability of the approach can be extended to multiple particle types (Fig. 2E).

### Subcellular mechanics during creep experiments with SEM^2^

While small-strain regimes are typical of a healthy physiology, we reasoned that high-strain situations occur in biology and are likely to produce more interesting subcellular mechanics^36^. Therefore, we performed another round of *in-silico* creep experiments for a fixed *N*_*p*_=1000 without (Fig. 3A) and with the nucleus (Fig. 3B, Supplementary Video SV2). To explore the large strain regime, we let the cell equilibrate for 10 s, stepped the applied stress to 20 Pa, and held it for 30 s before allowing it to relax for another 30 s (blue dashed line, Fig. 3C and D). We assessed the resulting cell strain (black solid line, Fig. 3C and D) and calculated the per-particle shear strain based on the reference configuration immediately before stress application (color-coded in Fig. 3A and 3B). Both with and without the nucleus, the strain increased with the applied stress and decreased upon stress removal. Notably, and differently from the small strain regime, the cell strain did not reach a steady state and was ∼2.5 times smaller in the presence of the nucleus (note that the reference configuration for the cell with the nucleus was 22% longer than without it). To analyze inter-particle distance statistically, we calculated the radial distribution function (rdf) (Figs. 3E and F). The rdf plot shows the relative abundance (Z-axis) of pairs of particles that are located at a given distance (X-axis) at various time points (Y-axis) during the simulation. In the equilibrium phase (dark green distributions, Fig. 3E and F), most particle pairs were near their equilibrium distance, *d*_*eq*_ ∼1.8 μm, as expected based on the rheology-preserving applied forces (Fig. 1B and Eqn. 1-2). Many particles were strained away from their equilibrium locations during the stretch (light green distributions, Fig. 3E-F). Finally, upon stress removal, particles could locally return to their equilibrium distance (grey distributions, Fig. 3E-F). Yet, due to the overall different configuration, the total number of particle pairs at *d*_*eq*_ decreased, providing a subcellular rationale for the plastic deformation observed at the cell level (Fig. 3C-D). These observations demonstrate that SEM^2^ can be applied to simulate and analyze cell-level and particle-level mechanics, thus offering a novel way to study subcellular mechanics in classical cell creep experiments.

**Figure 3:**
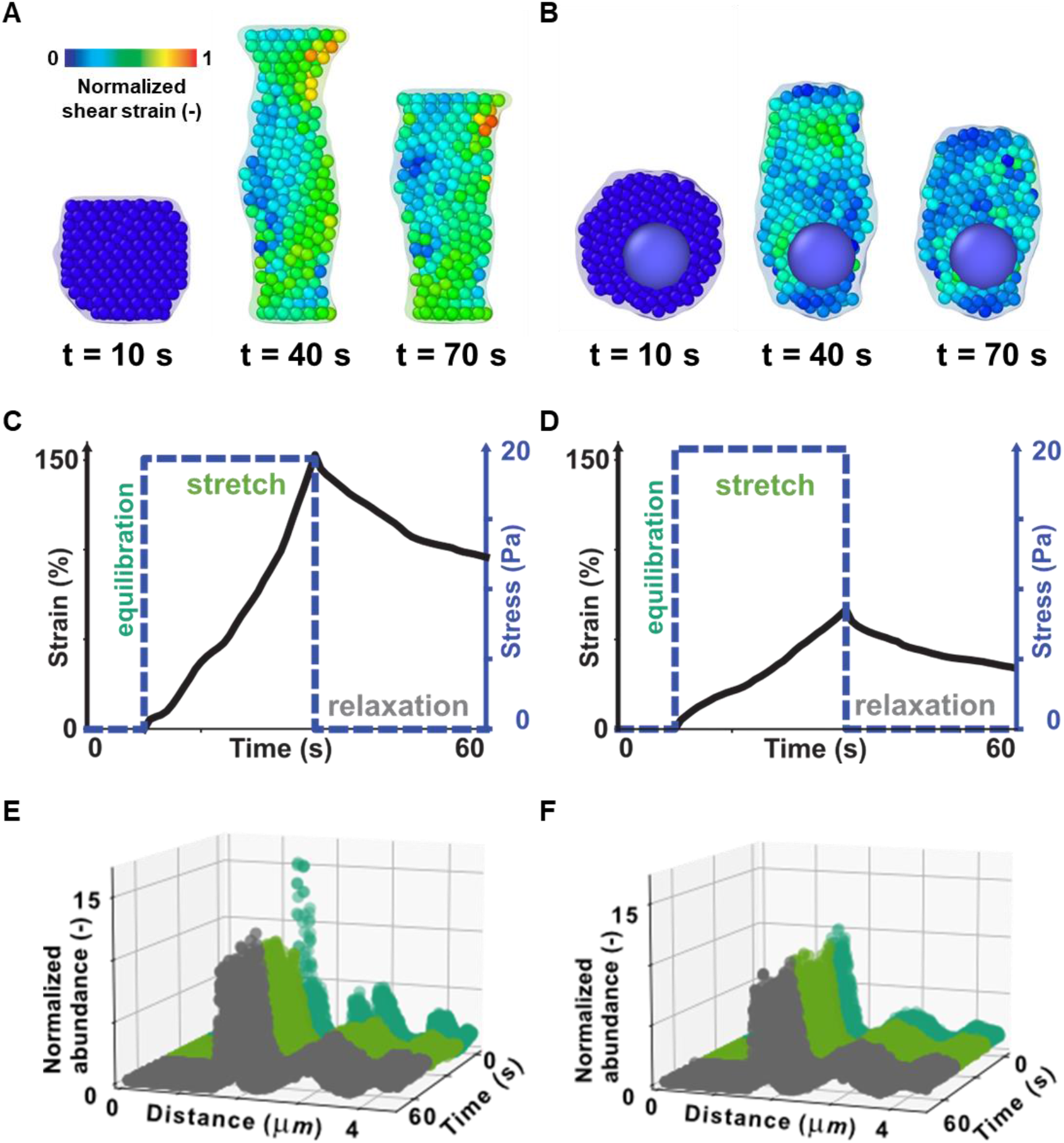
Subcellular mechanics with SEM^2^. (A-B) Representations of cells (*N*_*p*_ = 1000) without (A) and with (B) nuclear particles, as they undergo a high-stress (20 Pa) creep experiment. Cell membranes and particles (excluding the nucleus) have been colored based on the von-Mises shear strain invariant of the per-particle Green-Lagrangian strain tensor. The strain has been computed with respect to the unloaded configuration and the color coding was normalized to the simulation maximum to facilitate comparisons. Large and small strains are indicated by hot colors and cold colors, respectively. (C-D) The applied stress and the resultant axial strain are plotted for a single cell without (C) and with the nucleus (D). (E-F) The radial distribution function (rdf) of cytoplasmic particles in the cell without (E) and with the nucleus (F) stacked for various time points during the simulations (dark green, light green, and grey indicates rdf traces of time points in the equilibration, stretch, and relaxation phases, respectively).

### Modeling migration and active nuclear transport with SEM^2^

The creep and scaling validations are fundamental to the claim that SEM is a rheology-preserving framework for studying tissue morphogenesis. However, to actually model morphogenesis, we must also model cell migration and proliferation. In SEM++^27^, particle ensembles move as we add a particle on one side of the cell (leading edge) and remove a particle from the other (trailing edge, Supplementary Video SV3). Importantly, the force balance in Eq. 1 (Fig. 1B) generates rapid re-adjustments of particle positions and velocities in the ensemble that restore cell rheology. The particle addition/removal frequency determines how fast a cell moves, which is captured in the model parameter migration time *T*_*m*_^27^. When *T*_*m*_ is large with respect to the simulation duration (*T*_*sim*_), the number of particle addition events is small, and the cell moves a small distance from its initial position (and *vice-versa*). However, we noticed that in SEM++ the nucleus position within the cell changed with varying *T*_*m*_ and could drift close to the cell trailing edge (Fig. 4A). The fact that the nucleus moved very little during migration without bias in SEM++ is probably because the interaction between cytoplasmic and nuclear particles is also restricted by the cut-off, and it becomes net positive in the direction of migration only when the nucleus has drifted towards the cell edge. While this behavior is observed in certain diseased conditions^37^, most healthy cells exhibit a degree of active nuclear transport that is not captured by simply modeling passive rheology. Therefore, we extended the equation of motion to:

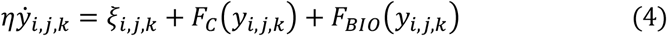

**Figure 4:**
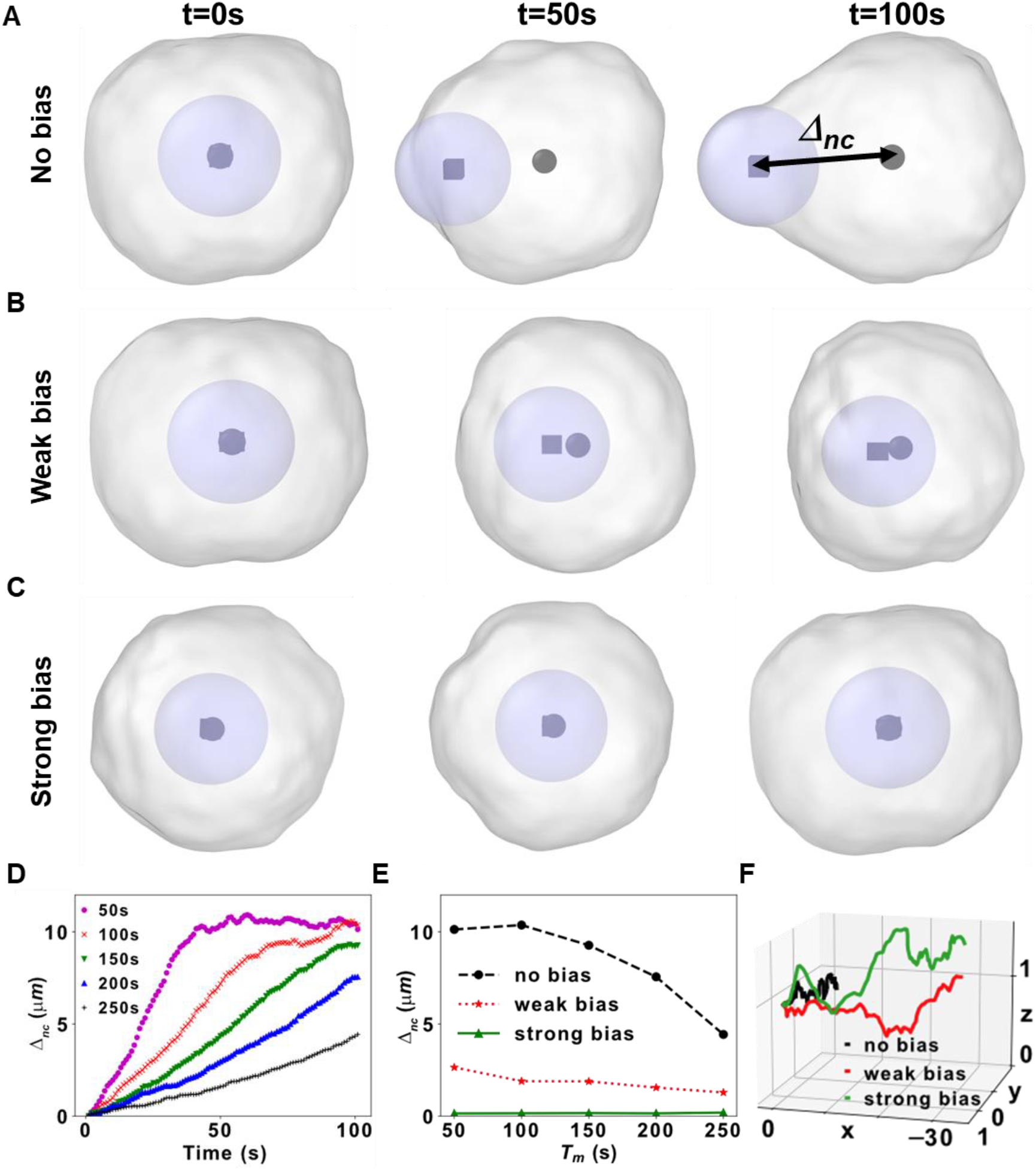
Nuclear transport and cell migration. (A-C) Cell migration snapshots at three different timepoints in the no-bias (A) weak bias, (B) and strong bias (C) conditions. The cell centroid is depicted as a black sphere, the nucleus centroid as a black cube. The distance between the nucleus and cell centers, Δ_*nc*_, is indicated in (A). (D) Δ_*nc*_ is plotted versus time with no bias for different values of *T*_*m*_. Note that the nuclear drift saturates at ∼10 um which is the radius of the cell. (E) Δ_*nc*_ versus *T*_*m*_ for the three different scenarios. (F) The full trajectories of the cell centers for the three different scenarios.

In Eq. (4), *F*_*BIO*_(*Y*_*i,j,k*_) is a general per-particle force and is intended to simulate accelerations imparted to certain particles by active biological mechanisms. In the case of active nuclear transport, we first introduced the variable Δ_*nc*_, which measures the distance between the centroid of the cell and the centroid of the nucleus; then, we specified *F*_*BIO*_ as a spring between these centroids (Fig. 4A, Methods). By adjusting the spring stiffness of this biasing mechanism, we could arbitrarily reduce Δ_*nc*_ to simulate various degrees of subcellular active transport ranging from it having no-bias (Fig. 4A), a weak bias (Fig. 4B), or a strong bias towards the cell centroid (Fig. 4C, also supplemental video SV4). For these simulations, we chose a *T*_*m*_ that would result in nuclear drift in the original SEM++ framework (*T*_*m*_ =100 s, *T*_*sim*_ = 100 s) and showed it being mitigated with different spring stiffnesses in SEM^2^. These simulations clearly showed that during migration, the centroids of each cell and its nucleus (depicted as a black sphere and black cube, respectively) would drift apart if not counteracted. To investigate how much nuclear drift affects cell migration, we performed more *in-silico* experiments altering *T*_*m*_ from 50 to 250 s while keeping the total duration to *T*_*sim*_ = 100 s. In the absence of *F*_*BIO*_, a degree of nuclear drift was observed for all *T*_*m*_ (Fig. 4D), but the dependence of Δ_*nc*_ on *T*_*m*_ was reduced or completely negated by increasing the nuclear biasing strength (Fig. 4E). Finally, the simulations showed that for the same choice of *T*_*m*_ and *T*_*sim*_, cells moved a smaller distance in the absence of biasing forces (see trajectories in Fig. 4F and Supplementary Video SV4), suggesting uncompensated nuclear drift affects cell motility.

### Subcellular mechanics during cell migration and nuclear transport with SEM^2^

To explain how the nucleus can slow down a cell without a biasing force (Fig. 4), we considered two particles, one near and another away from the nucleus. We observed that the two particles moved much farther apart in the no-bias condition than they did under strong bias (Fig. 5A). In fact, the analysis of the full trajectories of the two particles during migration with no bias (Fig. 5B) and with a strong bias (Fig. 5C) indicated that the particle near the nucleus barely moved in the direction of migration.

**Figure 5:**
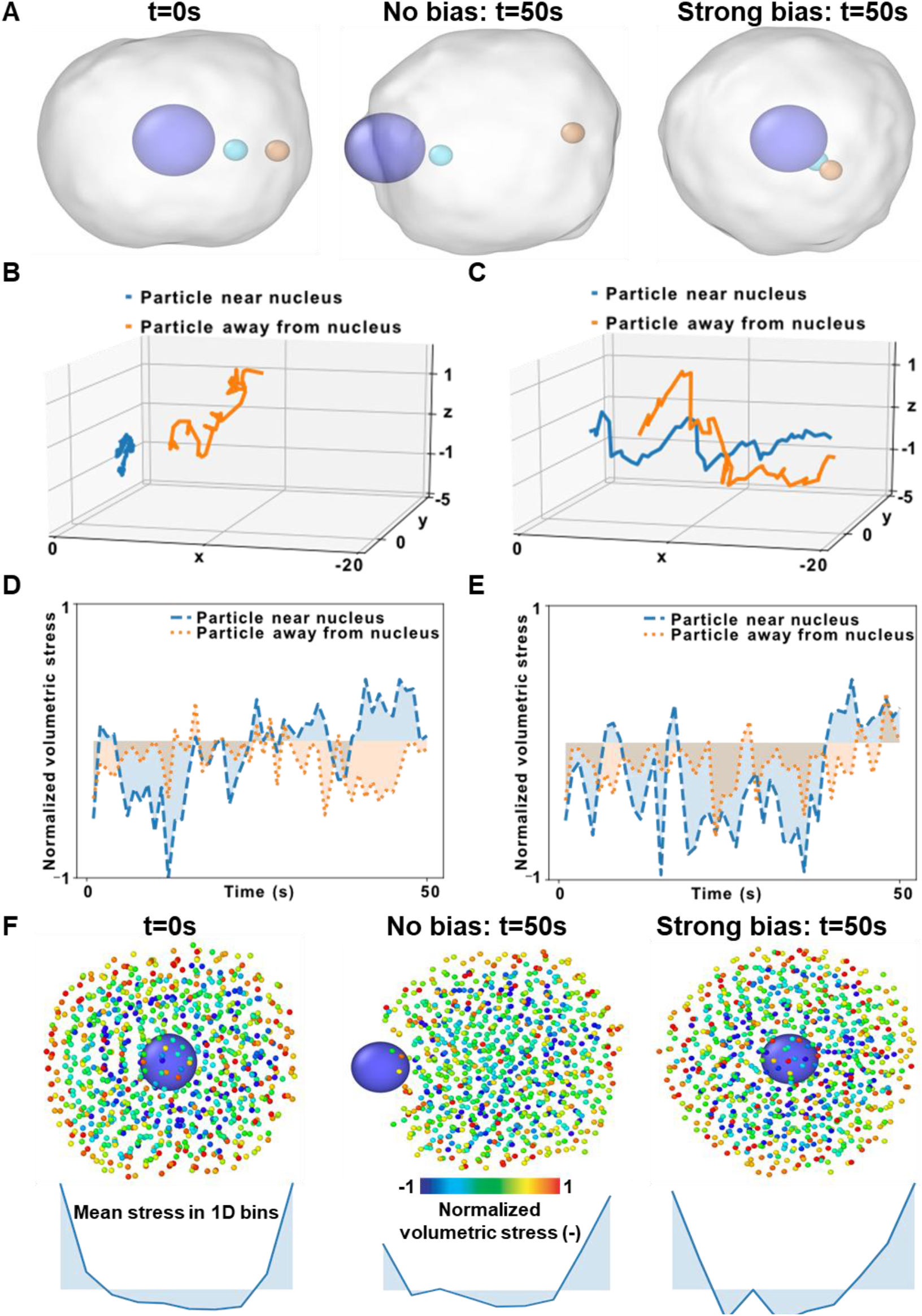
Subcellular mechanics during cell migration. (A) The locations of two particles, one near and another away from the nucleus, are shown before migration and after migrating for 50 s with no bias and strong bias. (B, C) The trajectories of the two particles during migration with no bias (B) and strong bias (C). (D, E) Normalized mean volumetric stress of a particle near the nucleus and another away from the nucleus during migration with no bias (D) and strong bias (E). (F) Cytoplasmic particles are color-coded per the normalized hydrostatic stress per particle in the cell before migration and after migrating for 50 s with no bias or strong bias. Mean particle stress in 1D bins is also plotted.

Next, we asked whether the potential between cytoplasmic and nuclear particles might also impact subcellular mechanics inside the whole cell. Particle-level strain (Fig 3) is ill-defined during migration simulations in SEM because particles are continuously added and removed from the ensemble, so there is no single reference particle configuration at the beginning of the experiment. Instead, we focused on per-particle mean volumetric stress, the relative extent of which can be assessed directly from the particle energy at any given time frame (see Methods). When we compared particles near the nucleus and away from it, we found that the magnitude and fluctuations of stress were higher in the particle near the nucleus for both the migration with no bias (Fig. 5D) and with a strong bias (Fig. 5E). However, in case of migration with a biasing force, the particle near the nucleus is mostly under greater stress (Fig. 5E) due to the push and pull of the nucleus, which is under a spring-like force towards the cell center.

We then asked whether this behavior was isolated to these two particles or existed across the cell. To answer this question, we color-coded the cytoplasmic particles of the migrating cells according to their stress normalized to the maximum stress computed during the simulation. Since the nucleus is a single rigid particle, we excluded it from this stress-dependent visualization (See Methods). We noticed that the mean volumetric stresses were mostly tensile (positive) along the cell membrane, whereas the inside was under compressive (negative) stress. To analyze the distribution of subcellular stress, we computed the average per-particle hydrostatic stress in 1D bins and found tensile peaks in correspondence with the cell membrane. For the case of migration with no bias, the nucleus is at a peripheral position resulting in an asymmetric stress distribution (Fig. 5F). However, for the case of migration with biasing force, since the nucleus is at the center of the cell, the particle stress distribution is roughly symmetric around the nucleus (Fig. 5F). Taken together, these results suggest that SEM^2^ can be used to model migration in more general cases than SEM++ and demonstrated how the per-particle analysis could be used to gain insights into subcellular mechanics during cell migration.

### Modeling cell proliferation in SEM^2^

Finally, to tackle problems in tissue morphogenesis with SEM, we also need to model cell proliferation. To implement proliferation, SEM++ considers two separate phases: Growth and division. In the growth phase, particles are added similarly to migration but without removal (See supplementary video SV5). When the particle number per cell doubles to 2*N*_*p*_, SEM++ enters the division mode. First, a second nuclear particle is created and allowed to interact with the original nuclear particle via a mildly repulsive potential, imitating how the poles of the mitotic spindle drift apart during cell division^38^. To nucleate a daughter cell, the new nuclear particle is given a new cell index (the k-th in the *Y*_*i,j,k*_ notation introduced in Eq. (1)). To complete cell division, the cell index of each cytoplasmic particle is re-assigned to one of the daughter cells based on the cell index of the closest nuclear particle, thus ensuring the splitting of the mother cell into two independent daughter cells. However, we preferred not to change the modeling assumptions regarding the nuclear particles to avoid artifacts due to particle size and weight changing during the simulation. Instead, SEM^2^ uses the nuclear-centering bias introduced in the previous section (see Methods). First, it creates the second nuclear particle close to the first at the center of the cell, thus modeling the increased DNA density observed before cell division^39^. Then it assigns cytoplasmic particles to the closer nucleus, thus splitting the cell at the center. Finally, the nuclear biasing mechanism recenters the nucleus of each daughter cell (Fig. 6A, Supplemental video SV6).

**Figure 6:**
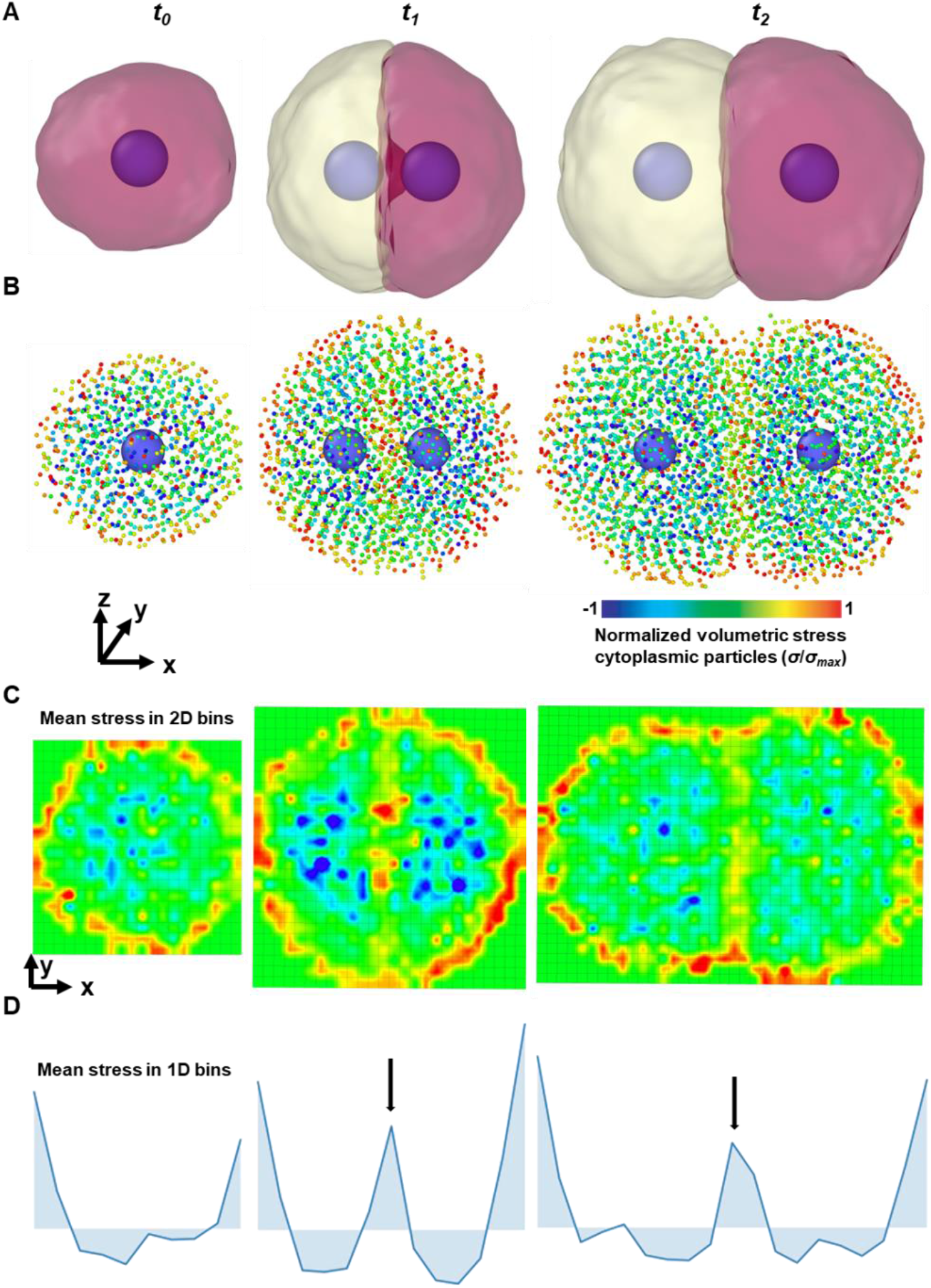
Cell proliferation. (A) Depiction of the cell proliferation simulation with nuclear particle and the membrane visualized with a Gaussian surface. (B) Depiction of the same proliferation simulations with cytoplasmic particles color-coded with normalized volumetric stress. The cell grows (from time, *t*_*0*_) with an increase in the number of particles, followed by duplication of the nucleus (*t*_*1*_), resulting in two daughter cells (*t*_*2*_). The interfaces between cells and the peripheral particles are under tensile (positive) stress, whereas the internal particles are under compressive (negative) stress. (C) Mean particle stress in 2D spatial bins on a projected x-y plane is color-coded. The cell membrane and the interface can be distinguished based on particle stress. Colormap is a per-particle in panel B and represents the average particle-level stress in the 2D bins in panel C. (D) Plot of mean per-particle hydrostatic stress in 1D spatial bins along the x-axis. Black arrows indicate the tensile peak corresponding to a cell-cell interface.

To study subcellular mechanics during cell division with SEM^2^, we resorted again to per-particle calculations and visualization by computing the mean volumetric stress per particle. Figure 6B shows a proliferating cell during growth and division with the cytoplasmic particles color-coded according to their normalized stress (see Methods). We noticed that the volumetric stresses are mostly tensile (positive) at the cell-cell boundaries and compressive (negative) inside. This could be better understood when we created spatial bins in two (Fig. 6C) and one (Fig. 6D) dimensions. In 2D, we observe tensile stresses at the outer rim of particle ensembles and the cell-cell interface (Fig. 6C), whereas in 1D we see tensile (positive) peaks (Fig. 6D arrows indicate peaks). Instead, particle volumetric stresses remained mostly compressive (negative) inside the cell (Figs. 6C and D). In other words, the gradient between compressive and tensile stress could be used to pinpoint the position of the cell membrane. This is particularly interesting because, so far in SEM, information about membranes has been prescribed with dedicated membrane particles or obtained algorithmically^27,32^.

### Modeling organoids and organ-chips with SEM^2^

To demonstrate that SEM^2^ can be used for tissue engineering, we investigated subcellular mechanics during cell proliferation in simulation scenarios mimicking organoids^1,4,5^ and organs-on-chips^6–8^. To do that, we performed three rounds of cell proliferation in unconstrained and constrained environments that model low-adhesion plates and the central section of a microfluidic channel, respectively. During the simulations, cells proliferated freely in the unconstrained environment (Fig. 7A) or along the channel in the constrained one (Fig. 7B, Supplementary Video SV7). We obtained the particle stress distribution in these two mechanical environments and depicted them in Figures 7C and 7D, respectively (Supplementary Video SV8). Again, particle stresses were tensile at the periphery and the cell-cell interfaces, whereas compressive inside. When we binned the particle stress along the unconstrained axis, the tensile peaks occurred when two or more cell-cell interfaces aligned, which was more frequent in the constrained environment (Fig. 7D).

**Figure 7:**
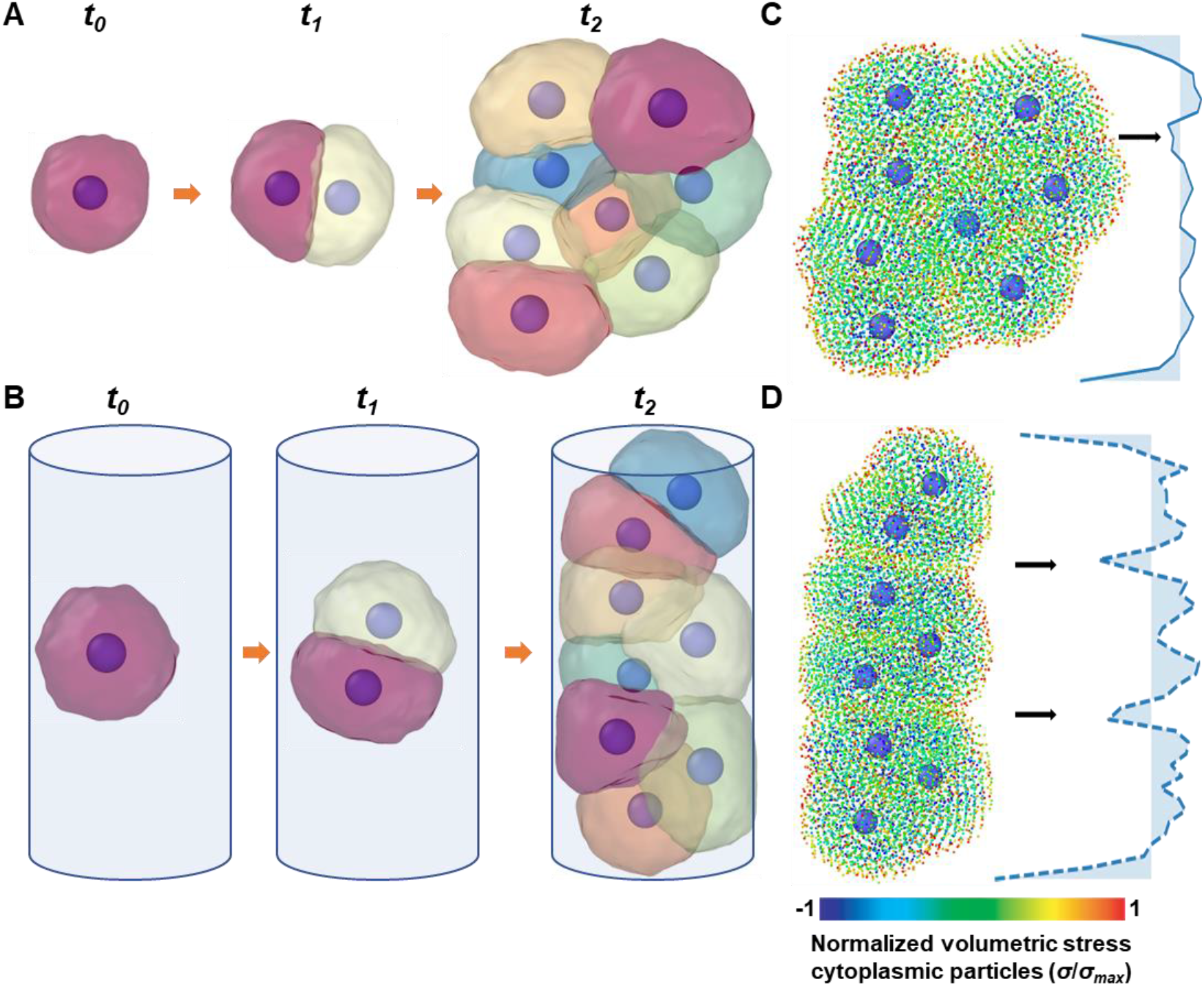
Traditional and engineered cell culture platforms. (A, B) Simulations of unconstrained (A) and constrained (B) proliferation simulations during which a single mother cell undergoes three rounds of cell division. (C,D) Color-coded particle stress distribution and binned mean particle stresses along the vertical axis are shown for unconstrained (C) and constrained (D) proliferation. Black arrows indicate the tensile peaks corresponding to the aligned cell-cell interfaces.

Finally, these simulations allowed us to look at the problem from the perspective of multiscale mechanics. At the tissue level, the volumetric stress versus time of the two tissues was the same (Fig. 8A), since it was dominated by cell proliferation unaffected by mechanical constraints. However, the axial strain was higher in the cylindrical channel (Fig. 8B), as expected from the mechanical environment that coerces newly formed cells to align axially (Fig. 7). Instead, at the cell level, volumetric strain followed the cell cycle: it increased as the cell grew and decreased when it divided in both constrained and unconstrained simulations (Fig. 8C). In other words, the growth of cells was not affected by the lateral constraint; instead, cells adapted their shape and orientation to make use of the unconstrained axial direction. This was confirmed by the particle-level stress distribution in 1D bins (Fig. 8D), where tensile peaks can be observed in correspondence of cell-cell interfaces, and they were higher and more frequent in the constrained condition (orange vs. cyan lines, black arrows in Fig. 8D). Together, these results suggest that SEM^2^ can be used to gain insights into multiscale and subcellular mechanisms of cell proliferation in traditional and engineered cell culture platforms.

**Figure 8:**
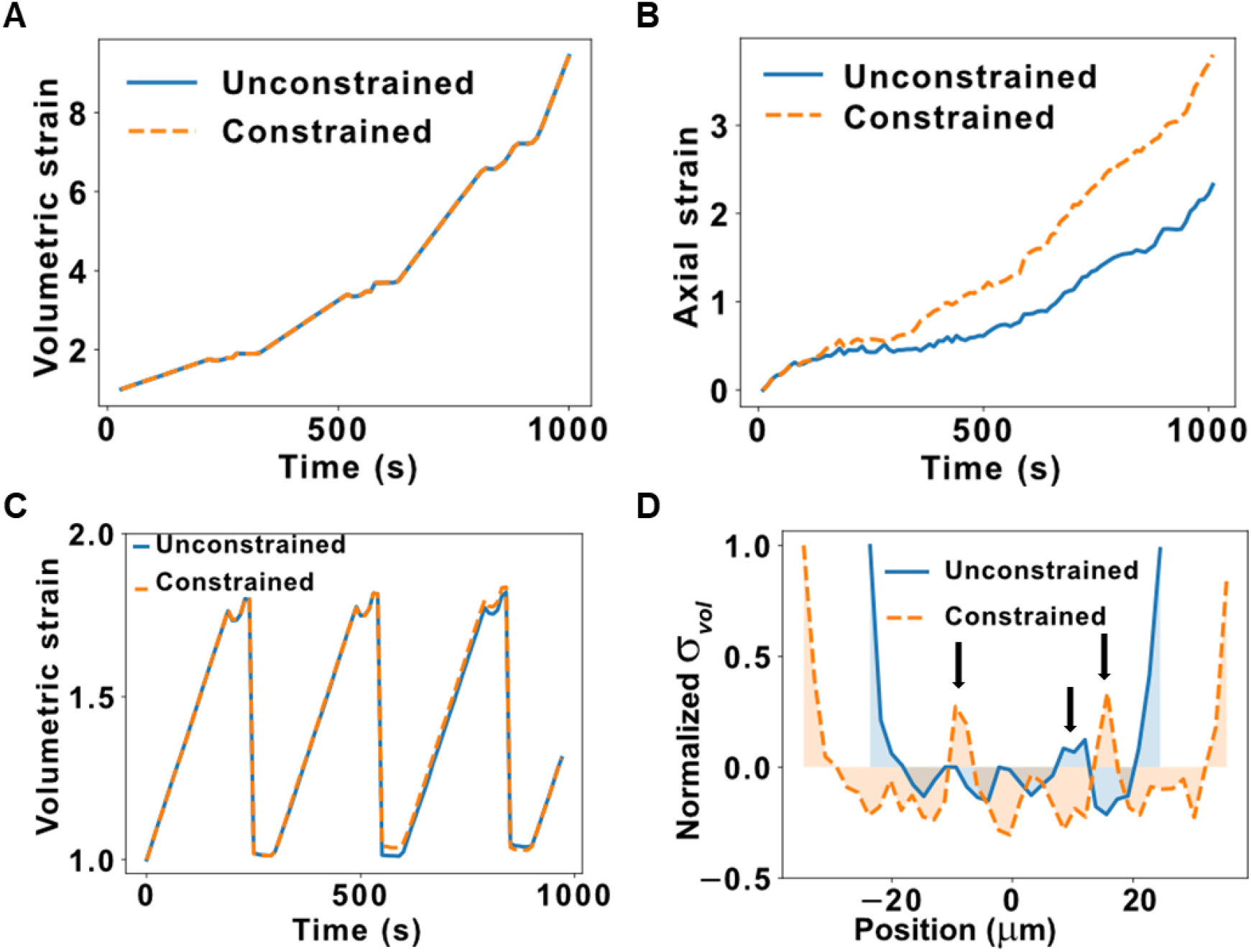
Stresses and strains in cell proliferation. (A) Unconstrained and constrained volumetric strain versus simulation time of the proliferating cells shown in Fig. 7. (B) Unconstrained and constrained axial strain along the unconstrained axis versus simulation time of the proliferating cells. (C) Unconstrained and constrained volumetric strain of a single cell versus time during the same proliferation simulation. (D) Mean particle stresses in 1D bins along the unconstrained axis for unconstrained and constrained proliferation. Black arrows indicate the tensile peaks corresponding to the aligned cell-cell interfaces.

## Discussion

We presented SEM^2^ as an updated version of SEM++ that enables the study of multiscale mechanics using coarse-grained particle dynamics. We extended the capability of SEM++ to model cell migration and proliferation^27^ by adding a general force term, *F*_*BIO*_, to the Langevin formulation (Eqn. 4). This force was used to model active nuclear positioning during migration (Fig. 4), as well as nuclear re-centering after cell division (Fig. 6). Furthermore, we demonstrated initial simulations of organoid- and organs-on-chips-like platforms using constrained and unconstrained proliferation (Fig. 7). Finally, in SEM^2^ we introduced particle-level strain (Fig 3) and stress calculations (Fig 5-7) downstream of the particle dynamics simulations, which enabled new statistical analyses and visualizations, such as the radial distribution function (Fig. 3) or the particle trajectories (Fig. 4-5)

Starting from a simple set of potentials derived from experimental cell stiffness and viscosity values in Eq. (3), SEM^2^ generated several interesting emergent mechanical behaviors consistent with recent experimental findings. For example, when exposed to high stress during creep experiments, cells deformed plastically in good agreement with many experimental observations, as reviewed recently^40^. Interestingly, we observed a ∼60% residual plastic deformation after high-stress creep (Fig 3C-D), which is in remarkable quantitative agreement with recent data from magnetic twisting cytometry where visco-elasto-plastic deformation of 3T3 fibroblasts^41^ was analyzed. Similarly, in cell migration simulations, our modeling work showed that nuclear positioning can affect the overall motility of the cells. When the nucleus lags at the trailing edge (our no-bias condition in Fig. 4), its positioning inside the cell is mediated by the pushing from the cell rear, as reviewed before^42,43^. Yet, we also saw that the cell accelerates when the force bias is used to push the nucleus actively toward the leading edge. This is consistent with recent evidence that cells relocate their nucleus towards the leading edge to overcome obstacles or constrictions^43,44^. During proliferation, we saw that the cell-cell interfaces are characterized by tensile stresses with opposite signs from the compressive stress observed inside the cell (Fig. 6-7). This behavior has also been studied experimentally, as many research groups have seen tensile forces at the cell-cell junction in-vitro^8,45^ and in-vivo during embryogenesis in zebrafish^46^. Finally, SEM^2^ predicted morphological re-arrangements of cell-cell interfaces to accommodate the same number of cells in constrained vs unconstrained simulations (Fig. 7-8). This observation is consistent with many experimental studies in engineered cell culture platforms, including hepatocytes in a liver-on-a-chip^47,48^ and cardiac muscle cells in 3D microfluidic^49^ and 2D micropatterned^50^ cell culture platforms. Importantly, in SEM^2^ these experimentally validated behaviors emerged spontaneously and were not imposed as modeling assumptions in all these examples. For example, tensile stresses measured at the cell-cell junctions are due to cortical actin dynamics, yet in SEM^2^, they emerge from interacting coarse-grained particles that do not explicitly represent actin or any intracellular component.

This study focuses on introducing the framework for subcellular mechanics to SEM++ and leveraged previous work for the choice of potential and particle types^23,27^. In the future, we envision using more heterogeneous subcellular particles to model various intracellular compartments (e.g., cytoskeleton, organelles). Similarly, we kept the SEM++ formalism and modeled the nucleus as a single heavy particle. However, this can be extended by adopting multiple nuclear particles. Finally, the subcellular mechanics approach we propose is agnostic to the choice of potentials^23^, which opens up the possibility of using different formulations, including data-driven approaches^51^. A major downside of particle-based methods is their computational cost. In SEM^2^, it took us ∼ 34 mins on four cores (AMD EPYC 7763, 2.45 GHZ) to perform the proliferation simulations with 8 cells and ∼10,000 particles. Yet, simulating a tissue consisting of ∼ 28000 cells with ∼ 33 million particles took ∼ 2 months (Supplementary Video SV9). Having built SEM++ and SEM^2^ on top of LAMMPS, we have a very scalable architecture for basic operations like updating the particles’ velocity field. However, every time we add or remove a particle from the particle list (e.g., proliferation), we introduce the intrinsically serial operation of updating the particle list, which increases overhead and decreases scalability. To reduce the impact of these limitations, dedicated parallelization schemes should be developed^52^.

In conclusion, we provide SEM^2^, an updated implementation of SEM++ that can be used to model subcellular mechanics during cell creep, migration, and proliferation experiments. The full code is available on GitHub (https://github.com/Synthetic-Physiology-Lab/sem2) for biologists, physicists, and engineers interested in modeling multiscale mechanics in shape-changing tissues.

## Methods

### Single-cell creep simulations

We have performed the simulations with LAMMPS, utilizing the package SEM++. We have created a cell consisting of *N*_*p*_ particles in a cylindrical region of space using the “fix sem_proliferate” functionality of SEM++. We have included walls on the top and bottom of the cell along the z axis, with “fix wall/lj93” functionality of LAMMPS. For example, we applied the 9-3 Lennard-Jones potential between walls and particles to model the adhesion of particles with the fixed and movable plates in the experiments. Additionally, there is a cylindrical constraint around the z-axis using the “fix indent” functionality of LAMMPS. We have employed Brownian dynamics time integration to update the position and velocity of particles, using the “fix bd” functionality of SEM++ along with the “fix langevin” functionality of LAMMPS. Upper and lower slabs of particles have been created considering all particles in the geometric regions of thickness 1.5 microns (∼ 10% of the cell height) at the cell top and bottom.

To make the bottom slab fixed, the velocity of the particles of this layer was always fixed as 0. The input stress, applied to the top layer, has three different segments: 1) hold, 2) load, and 3) unload, as shown in Fig. 2. The stress,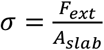, applied to the top layer is 0, σ, and 0, respectively (Fig. 2). The force to be applied on each particle during the loading segment is computed as 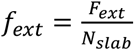 where _*Nslab*_ is the number of particles of the movable slab, as depicted in Fig. 2. To quantify the slab surface area for computing the applied stress we generated a surface mesh on selected subcellular particles, employing the Gaussian density method in OVITO^53^. To obtain the total contact surface area, we then summed all triangular mesh facets, in which all three vertices have stretch direction coordinate values more than a threshold. The threshold depends on the cell shape and is chosen such that the resultant surface area encompasses all surface particles in the movable slab. The nominal stress can then be computed as the ratio of the total force acting on the slab and the contact surface area. The axial strain is calculated as 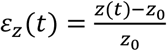, where *z*_0_ is the initial height of the cell before the load segment, and z(*t*) is the height at time *t*. Two more samples have been generated by uniformly rotating the sample around the x-axis.

We calculated the per-particle strain tensor in OVITO using the “atomic strain” modifier. The atomic strain computation in OVITO is based on the finite strain theory. The Green-Lagrangian strain tensor, **E**, is calculated from the deformation gradient tensor of each particle. The von-Mises shear strain invariant, *γ*, for each particle, is then computed as

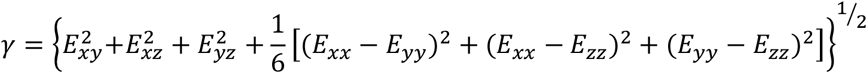

We computed the strain tensor with reference to the initial configuration before loading, i.e. at *t*=10s, with a cutoff radius of 10 μm. For visualization in Figures 3A and 3B, we have normalized all per-particle strains with the highest magnitude of shear strain observed across all simulations.

### Migration and proliferation

We created a cell of *N*_*p*_ particles in free space using the “fix sem_PMN” functionality of SEM++ without the walls perpendicular to the z-axis or the cylindrical shell centered around the z axis. We implemented migration with the “fix sem_PMN” functionality of SEM++. This is a polarity-induced migration, where the direction of migration depends on the shape of the cell and is computed based on the polarity of the cell.^27^ A *migration event* occurs when particles are added near the leading edge and removed from the trailing edge. *Migration events* can be observed in Supplementary Video SV3. The probability of a *migration event* is inversely proportional to the parameter migration time, *T*_*m*_, and proportional to *N*_*p*_. In between *migration events*, Brownian Dynamics time integration is carried out using the “fix bd” functionality of SEM++ and the “fix langevin” functionality of LAMMPS. Similarly, a *proliferation event* occurs when particles are added^27^, as shown in Supplementary Video SV5. Proliferation events are directly proportional to *N*_*p*_ and inversely proportional to the total proliferation time. When the number of particles in a cell reaches 2*N*_*p*_, the nucleus is replicated as explained in the main text (Supplementary Video SV6).

To model active transport of the nucleus during migration, we incorporated a biasing force, which acts like a spring force between the cell center of mass (COM) and the nucleus. At each timestep, the distance between the cell center of mass and nucleus, *Δ*_*nc*_, is computed. We utilized the “fix addforce” functionality of LAMMPS to incorporate a force on the nuclear particle proportional to the square of the distance, *Δ*_*nc*_. The constant of proportionality can be tuned to model nuclear position in various cell types, ranging from those in which the nucleus is at the cell center to those close to the cell periphery. We have incorporated this functionality of adding a *Δ*_*nc*_-dependent force bias fully from the LAMMPS input script, without necessitating any change in the source code.

For biasing the nucleus towards the center, let us consider the equation of motion after Newton’s law:

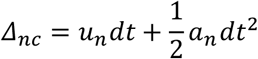

In the absence of other forces, the nucleus would reach the *current* center of mass of the cell in one timestep, *dt*, if *u*_*n*_ and *a*_*n*_ are the velocity and acceleration of the nuclear particle, respectively. If *K* is the elastic constant of the spring connecting the nucleus to the cell center, the biasing force on the nucleus is defined by

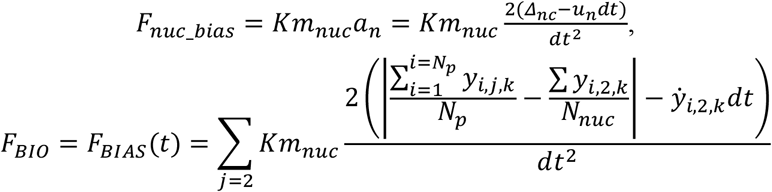

### Per particle stress visualizations

The per-particle stress has been determined utilizing the “compute stress/atom” functionality of LAMMPS. This yields per-particle virial stress, which has units of energy. Since this is an overdamped system, the contribution to energy and stress is primarily from potential energy. The stress tensor for atom *I* is defined as

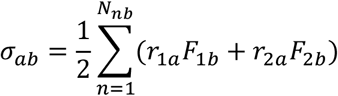

where σ_*ab*_ is the stress; *a* and *b* take on values from *x, y, z* to generate components of the tensor *N*_*nb*_ is the number of neighbors of the i-th atom. *r*_1_, *r*_2_, *F*_1_, *F*_2_, are positions and forces due to pairwise interactions.

We computed each particle’s mean volumetric stress (in energy units) as 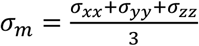. To distinctly visualize the distribution of stresses across the subcellular material, we chose a look-up table that emphasizes contrast for most computed values. We provided an analysis of this in Supplementary Figure S2, and the particle stress distribution of a migrating cell can be seen in Figure 5F. It should be noted that in Fig. 5F, the radius of the cytoplasmic particles is smaller than *d*_*eq*_/2 to visualize the stress distribution among all particles inside the cell.

We utilized the “spatial binning” modifier in OVITO to further analyze the spatial distribution of stresses. We computed the mean particle stress inside 1-D bins and normalized it by the maximum magnitude of stress across all bins and plotted it in Figure 5F.

## Data availability statement

The data that supports the findings of this study are available within the article and its supplementary material.

## Ethics approval statement

Not required.

