## Supplementary figures and video captions for "SEM^2^: A computational framework to model multiscale mechanics with subcellular elements"

* co-corresponding authors

#### Abstract:

Modeling multiscale mechanics in shape-shifting biological tissues in embryos, traditional, or engineered cell culture platforms (organoids, organs-on-chips) is both important and challenging. In fact, it is difficult to model relevant tissue-level structural changes mediated by discrete events at the cellular and subcellular levels, such as migration and proliferation. To accomplish this, we leveraged the subcellular element modeling (SEM) method, where ensembles of coarse-grained particles interacting via empirically defined potentials are used to model individual cells while preserving cell rheology. However, an explicit treatment of multiscale mechanics in SEM was missing. Here, we introduced SEM$^2$, an extended version of the open-source software SEM++ and LAMMPS, enabling new analyses and visualization of particle-level stress and strain. We demonstrated various functionalities of SEM$^2$ by simulating cell creep, migration, and proliferation in scenarios that recapitulate classical and engineered cell culture platforms. For every scenario, we highlight key mechanobiology that emerges spontaneously from particle interactions and discuss recent experimental evidence as qualitative validations of our simulations. The code for SEM2 is available on GitHub at https://github.com/Synthetic-Physiology-Lab/sem2 .

### Supplementary figures

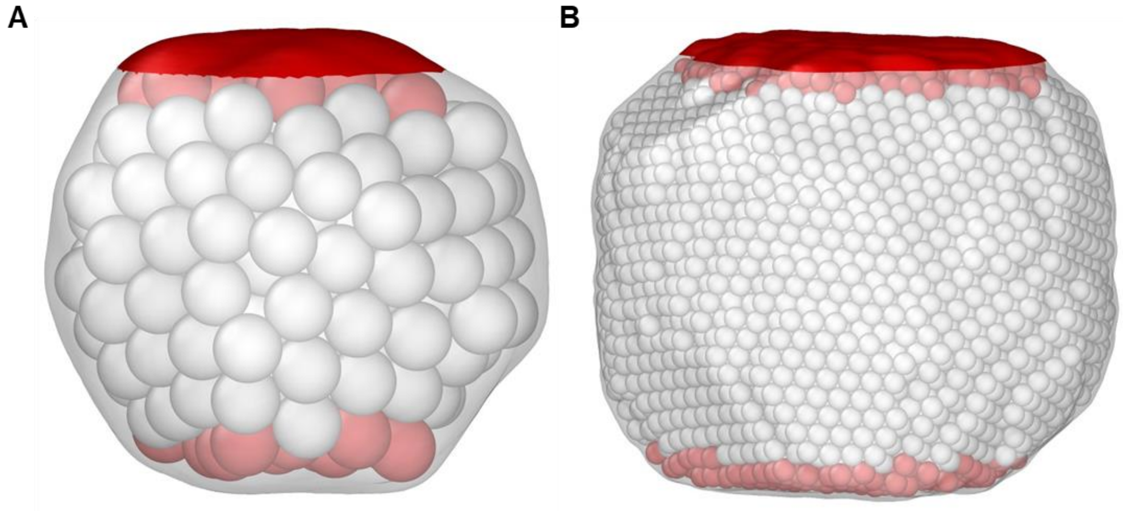


**Figure S1. Single cell creep: illustration of pre-stretched configuration.** (A) A cell in the pre-stretched configuration is plotted using a Gaussian surface mesh (white) that encloses all particles for $N_p$=250. The cell is sandwiched between a fixed and a movable plate. The fixed and movable particles are red whereas the other particles are white. The top's red (shaded) surface indicates the area considered for stress calculation. (B) A cell with $N_p$=10000 in the pre-stretched configuration.

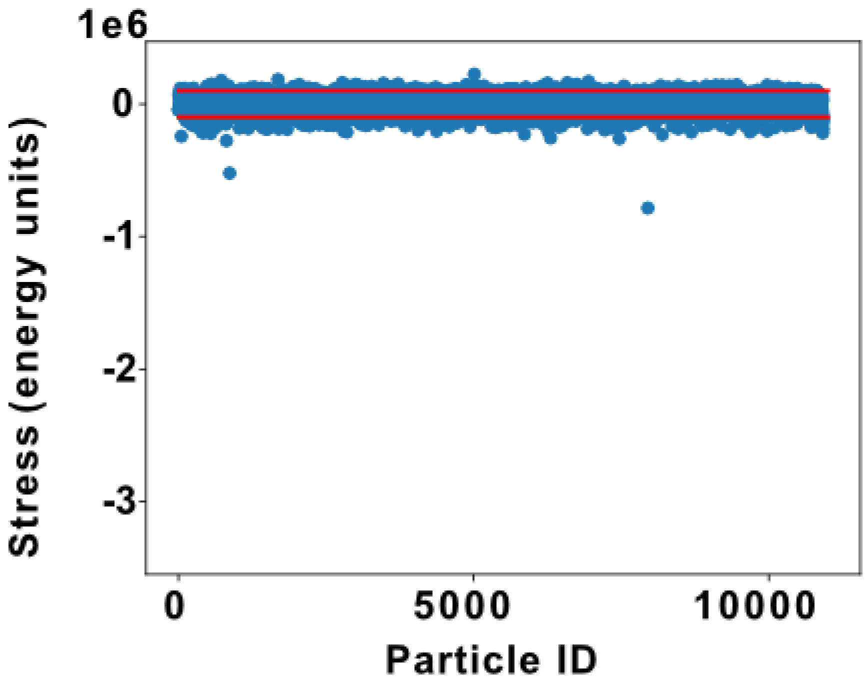


**Figure S2**: Particle stresses for all particles during the proliferation simulation. The stresses of majority of the particles lie in the range (-100000, 100000) depicted by the red lines. Thus, this range was used generate a look-up table that was used in the portrayal of particle stresses in Figs. 5F, 6B, 7C and 7D. For example, stresses within the above range were normalized by 100000 to obtain a number between -1 to 1 and was assigned a color according to the color map shown in the respective figures. All stresses greater than 100000 were assigned the hottest color and those less than -100000 were assigned the coldest color.

### Supplementary video captions

**Supplementary video 1:** Small stress creep experiment with cells without (left) and with (right) the nucleus.

**Supplementary video 2:** Large stress creep experiment with cells without (left) and with (right) the nucleus.

**Supplementary video 3:** Particle polymerization (green) and depolymerization (blue) during cell migration

**Supplementary video 4:** Migrating cells with no-, weak-, and strong-nuclear biasing mechanisms.

**Supplementary video 5:** Particle polymerization (green) during one cell division

**Supplementary video 6:** One cell division with various visualization of the cell (top row) and the subcellular particles (bottom row). Membranes are color-coded by cell id (top left) or by normalized stress at the periphery (two points of view, top right). Particles color-coded by particle-level stress in the 3D volume (bottom left, smaller particle visualization), or on two vertical slices (bottom right).

**Supplementary video 7:** Cell proliferation in unconstrained (left) and constrained (right) environments mimicking organoids and microfluidic channels, respectively.

**Supplementary video 8:** Cell proliferation in unconstrained (left) and constrained (right) environments, color-coded as described in supplementary video 6.

**Supplementary video 9:** Proliferation simulation resulting in a tissue consisting of ~ 28000 cells with ~ 33 million particles. Particles were colored according to their cell number.
